# Tn5 transposase-based epigenomic profiling methods are prone to open chromatin bias

**DOI:** 10.1101/2021.07.09.451758

**Authors:** Meng Wang, Yi Zhang

## Abstract

Epigenetic studies of rare biological samples like mammalian oocytes and preimplantation embryos require low input or even single cell epigenomic profiling methods. To reduce sample loss and avoid inefficient immunoprecipitation, several chromatin immuno-cleavage-based methods using Tn5 transposase fused with Protein A/G have been developed to profile histone modifications and transcription factor bindings using small number of cells. The Tn5 transposase-based epigenomic profiling methods are featured with simple library construction steps in the same tube, by taking advantage of Tn5 transposase’s capability of simultaneous DNA fragmentation and adaptor ligation. However, the Tn5 transposase prefers to cut open chromatin regions. Our comparative analysis shows that Tn5 transposase-based profiling methods are prone to open chromatin bias. The high false positive signals due to biased cleavage in open chromatin could cause misinterpretation of signal distributions and dynamics. Rigorous validation is needed when employing and interpreting results from Tn5 transposase-based epigenomic profiling methods.

## Introduction

Due to the sample loss and inefficient immunoprecipitation of traditional chromatin immunoprecipitation (ChIP)-based methods, low-input epigenomic profiling methods are needed for studying rare samples such as mammalian oocytes and preimplantation embryos^1^. Several low-input chromatin immunoprecipitation followed by sequencing (ChIP-seq) methods including ULI-NChIP^2^, scChIP-seq^3^ and STAR ChIP-seq^4^ have been developed. To overcome inefficient immunoprecipitation, immunoprecipitation-free methods, such as CUT&RUN^5^ and scChIC-seq^6^ that use chromatin immuno-cleavage (ChIC)^7^ strategy, have been developed. These low-input methods have been widely used in studying epigenome reprogramming during early embryonic development which have revealed distinct dynamics of different epigenetic markers^4, 8–12^.

Recently, the Tn5 transposase-based^13^ library construction is getting popular because it can fragment DNA while simultaneously adding library adaptors thus simplifying experimental procedures and reducing sample loss^14, 15^. Tn5 transposase-based epigenomic profiling methods utilize Protein A (or Protein G) fused with Tn5 transposase (pA-Tn5) to cleave DNA at the targets of primary antibody, allowing all procedures to be completed in the same tube without immunoprecipitation step, which largely avoided sample loss. Several Tn5-based methods have been developed to capture histone modifications or transcription factor (TF) binding using small number of cells or even single cell, including CUT&Tag^16^, CoBATCH^17^, ACT-seq^18^, itChIP-seq^19^, ChIL-seq^20^ and Stacc-seq^21^. However, the Tn5 transposase is known to prefer accessible DNA regions^22^. It has been noted that some of the Tn5-based methods are confounded by DNA accessibility^23^, but no systematic comparative analysis has been done to determine to what extent the results of these methods are affected by chromatin accessibility. Here we present systematic comparative analyses which reveal that, for some of the methods, overall ~30-50% false positive peaks can be contributed by open chromatin artefacts. Such high level of false positive peaks could affect data interpretation leading to false conclusions, which raises concerns on choosing, developing and interpreting results from Tn5-based epigenomic profiling methods.

## Results

### Tn5-based epigenomic profiling methods have varied level of biases toward open chromatins

CoBATCH^17^, CUT&Tag^16^, ACT-seq^18^ and Stacc-seq^21^ are very similar methods, which are based on the in situ immuno-cleavage strategy (**Fig. 1a**). CoBATCH and CUT&Tag add primary antibodies first, then add the pA-Tn5, while ACT-seq and Stacc-seq pre-incubate primary antibodies with pA-Tn5. However, besides cleavage at the target sites, the free pA-Tn5 has the potential to tagment open chromatins. To wash out free pA-Tn5, these methods employ different washing conditions. The CUT&Tag uses high salt (300 mM NaCl) washing to suppress background tagmentation, CoBATCH and ACT-seq utilize milder washing conditions, while the washing step is optional in Stacc-seq. The itChIP-seq, on the other hand, is an immunoprecipitation-based method that utilizes Tn5 to tagment DNA first before antibodies are added to pull-down the target fragments. Thus, theoretically the itChIP-seq method should not be confounded by open chromatins as antibody-based selectivity is applied after tagmentation.

**Fig. 1.**
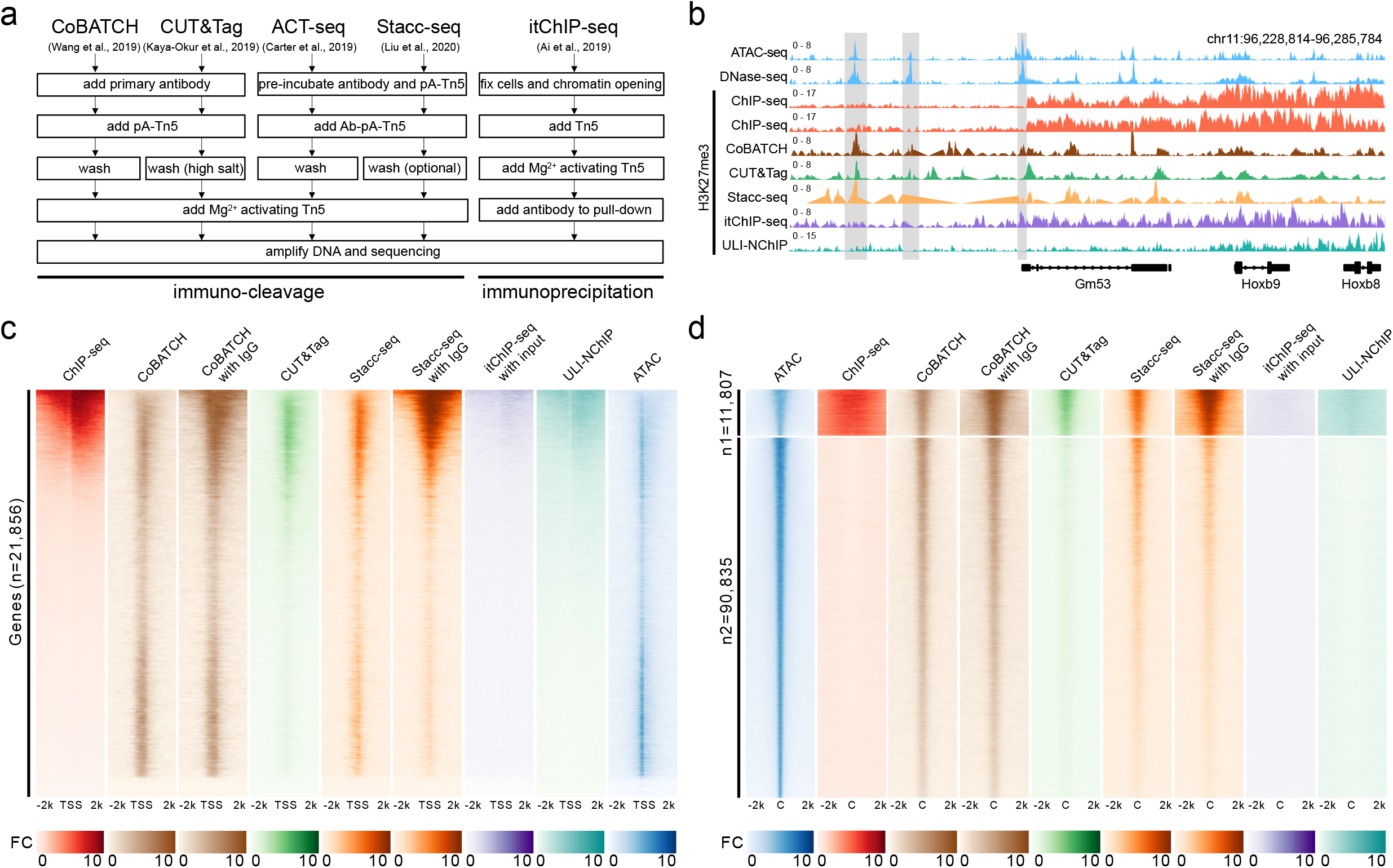
Signal distributions of Tn5-based epigenomic profiling methods. **a**, Major experimental procedures for different Tn5-based epigenomic profiling methods. pA-Tn5: Protein A and Tn5 fusion complex; Ab: primary antibody. **b**, Genome browser snapshot around Hoxb cluster in mESCs for open chromatin fold-change signals (ATAC-seq and DNase-seq), and H3K27me3 fold-change signals for two ChIP-seq datasets, Tn5-based methods and ULI-NChIP. The itChIP-seq fold-change signals were normalized by input. The signals for CoBATCH, CUT&Tag and Stacc-seq were fold-changes over background. **c**, H3K27me3 signal enrichment for different methods around the transcription start sites (TSS±2kb) of mouse coding genes, and was compared to ATAC-seq signals around the TSSs (FC: fold-change over background/input). **d**, Signal enrichment for different methods at all open chromatin regions (n1: open chromatin regions with H3K27me3 ChIP-seq signals; n2: open chromatin regions without H3K27me3 ChIP-seq signals; C: center of ATAC-seq peaks; FC: fold-change over background/input).

To perform systematic comparative analysis of the Tn5-based epigenomic profiling methods, we collected publicly available H3K27me3 data (**Supplementary Table 1**) of mouse embryonic stem cells (mESCs) generated by CoBATCH, CUT&Tag, Stacc-seq and itChIP-seq. Since the protocols for ACT-seq and Stacc-seq are almost identical, and the original ACT-seq study did not include H3K27me3, we analyzed the Stacc-seq data with conclusions applicable to ACT-seq. The bulk ChIP-seq of H3K27me3 in mESCs was used as a reference for comparing the H3K27me3 peaks derived from these Tn5-based methods. The two different bulk ChIP-seq datasets^24, 25^ were highly similar (**Supplementary Fig. 1a, b**) and the peaks were considered as true positive peaks in mESCs. To determine whether each Tn5-based method was confounded by open chromatin, we asked whether peaks that were not overlapped with bulk ChIP-seq peaks were instead overlapped with open chromatin peaks derived from ATAC-seq in mESCs. The open chromatin revealed by ATAC-seq^26^ were highly similar to that revealed by DNase-seq^25^ in mESCs (**Supplementary Fig. 1c, d**). As a control for low-input epigenomic profiling method without using Tn5 transposase, we included the H3K27me3 dataset in mESCs generated by ULI-NChIP^8^. A genome browser view around the Hoxb locus comparing the signals of Tn5-based methods with those of bulk ChIP-seq, ULI-NChIP, and open chromatin (ATAC-seq and DNase-seq) (**Fig. 1b**) revealed: 1) CoBATCH, CUT&Tag, and Stacc-seq detected H3K27me3 peaks not present in ChIP-seq or ULI-NChIP but overlapping with ATAC-seq and DNase-seq peaks (shaded); 2) For peaks overlapping with ChIP-seq, the peak patterns were more similar to ATAC-seq and DNase-seq rather than the ChIP-seq; 3) itChIP-seq showed the most similar pattern to that of the ChIP-seq in this region, which was coincident with the fact that the immunoprecipitation-based itChIP-seq procedure is different from the other pA-Tn5 immuno-cleavage-based methods (**Fig. 1a**).

The above observation raised the possibility that at least some of the Tn5-based methods may be biased toward open chromatin to generate false positive peaks. To explore this possibility, we analyzed the overall signal distribution of these methods by first focusing on the transcription start sites (TSS) of all coding genes. An analysis of the ChIP-seq datasets indicated that the H3K27me3 signals were enriched in the TSSs of a subset of genes consisted of mainly the Polycomb-group (PcG) targets, but with the majority of the genes, mostly of non-PcG targets, lack the H3K27me3 signals around their TSSs (**Fig. 1c**). However, the CoBATCH and Stacc-seq methods detected H3K27me3 enrichment at almost all the TSS regions including the non-PcG targets, which were more similar to the open chromatin patterns detected by ATAC-seq (**Fig. 1c**). The CUT&Tag method detected a weak signal enrichment at the TSSs without H3K27me3 ChIP-seq signals. The itChIP-seq and ULI-NChIP methods detected a pattern more similar to that of ChIP-seq although their signals were generally weaker (**Fig. 1c**). itChIP-seq is not an immunoprecipitation-free method, thus its signals need to be normalized by input control, similar to ChIP-seq and ULI-NChIP. For the immunoprecipitation-free methods, input DNA or IgG control is usually not needed for signal normalization. Nevertheless, we tested whether the confounding of open chromatin signals could be eliminated by using input/IgG control. Using the publicly available input/IgG controls for CoBATCH and Stacc-seq (no input/IgG control for CUT&Tag), we recalculated the H2K27me3 enrichment and found that normalizing with input/IgG did not improve the CoBATCH results (**Fig. 1c**). On the other hand, this normalization did enhance the signals of Stacc-seq overlapping with ChIP-seq, while reduced the signals not overlapping with ChIP-seq (**Fig. 1c**). However, the IgG control normalized Stacc-seq H3K27me3 profile was still more similar to the ATAC-seq profile than that of the H3K27me3 ChIP-seq at the non-PcG targets (**Fig. 1c**).

Next, we focused our analysis on open chromatin regions by dividing the open chromatin regions into two groups that with or without H3K27me3 ChIP-seq signals (**Fig. 1d**). The CoBATCH and Stacc-seq detected signals exhibit a clear enrichment at the open chromatin regions without ChIP-seq signals, and input/IgG control normalization did not change the situation. The CUT&Tag method detected weak signals, while itChIP-seq and ULI-NChIP detected no signals at the open chromatin regions without ChIP-seq signals (**Fig. 1d**). These results indicate that some of the Tn5-based methods, particularly the CoBATCH and Stacc-seq, are biased toward open chromatin peaks that do not have H3K27me3 ChIP-seq signals.

### Open chromatin is the source of false positive peaks detected by Tn5-based methods

Next we performed quantitative analysis to determine the level that each of the Tn5-based method is confounded by open chromatin. To this end, we used the same criteria (p-value < 1e-4 and q-value < 0.01) in peak calling for each method. Peaks that overlapped with ChIP-seq peaks were considered as true positives. Peaks that did not overlap with ChIP-seq peaks were considered as potential false positive signals, and were further analyzed to determine whether they could be mapped to open chromatin (**Fig. 2**). We found 5,189 out of the 9,125 CoBATCH peaks did not overlap with ChIP-seq peaks, but were mapped to open chromatin with strong correlation to ATAC-seq signals (**Fig. 2a**). Indeed, 82.3% (4,270 out of 5,189) of the CoBATCH peaks that did not overlap with ChIP-seq peaks were overlapped with ATAC-seq peaks (**Fig. 2b**). Peaks derived from CoBATCH normalized by IgG control showed even more false positives and ATAC-seq signals also enriched in these false positive peaks (**Supplementary Fig. 2a**). A similar analysis of the CUT&Tag dataset revealed 1,387 out of the 6,805 peaks were not overlapped with ChIP-seq peaks (**Fig. 2c**). Of these non-overlapping peaks, 24.2% (335 out of 1,387) were overlapped with ATAC-seq peaks (**Fig. 2d**). For Stacc-seq, 1,687 out of the 6,190 peaks were not overlapped with ChIP-seq peaks, but showed strong open chromatin signals (**Fig. 2e**). Of these non-overlapping peaks, 75.6% (1,275 out of 1,687) were overlapped with ATAC-seq peaks (**Fig. 2f**). Peaks derived from Stacc-seq normalized by IgG control still showed open chromatin enrichment for the false positive peaks (**Supplementary Fig. 2b**). On the other hand, the itChIP-seq peaks that showed no overlap with ChIP-seq peaks also did not overlap with ATAC-seq peaks (**Fig. 2g, h**, **Supplementary Fig. 2c**). For ULI-NChIP, although about half of the peak regions showed no ChIP-seq signals, no ATAC-seq signals were detected in these non-overlapping regions (**Fig. 2i, j**). Collectively, these results indicate that the great majority of the non-overlapping peaks detected by CoBATCH or Stacc-seq, and some of the non-overlapping peaks detected by CUT&Tag are mapped to open chromatin regions, and they could be false positive artefacts.

**Fig. 2.**
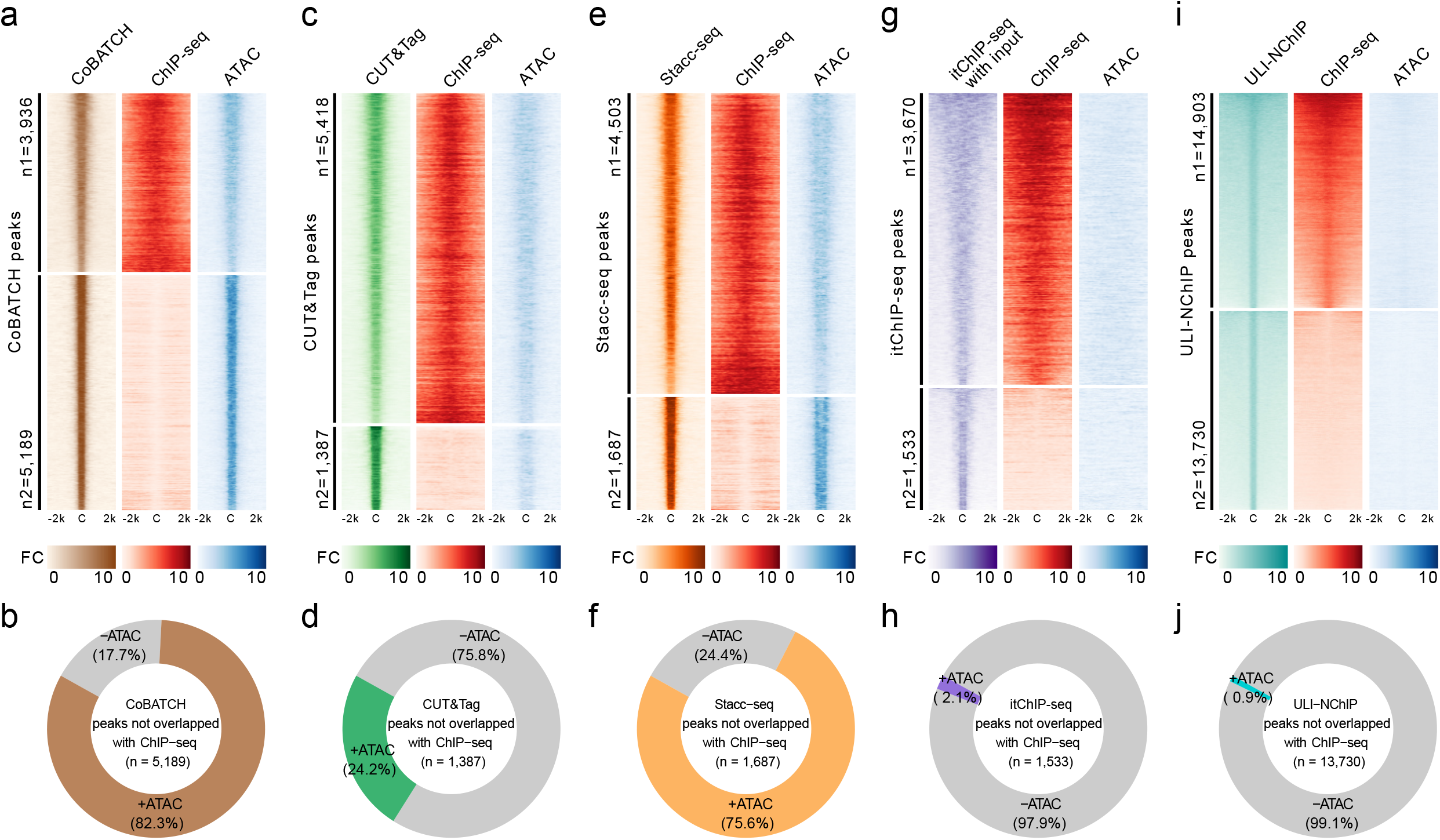
Evaluation of peaks from Tn5-based epigenomic profiling methods. Significant peaks (p-value<1e-4 and q-value<0.01) called from each method (**a**: CoBATCH, **c**: CUT&Tag, **e**: Stacc-seq, **g**: itChIP-seq, **i**: ULI-NChIP) were divided into two parts: n1 – peaks overlapping with ChIP-seq peaks, n2 – peaks not overlapping with ChIP-seq peaks, and compared with open chromatin signals measured by ATAC-seq (C: center of peaks called from each method; FC: fold-change over background/input). The itChIP-seq results shown in **g** and **h** were from 10k cells. For each method (**b**: CoBATCH, **d**: CUT&Tag, **f**: Stacc-seq, **h**: itChIP-seq, **j**: ULI-NChIP), peaks that were not overlapped with ChIP-seq peaks were further compared to ATAC-seq peaks (+ATAC: overlapping with ATAC-seq peaks; −ATAC: not overlapping with ATAC-seq peaks).

### High false positive rate due to open chromatin affected global distribution of peaks

To determine the relative reliability of the different Tn5-based epigenomic profiling methods, we next calculated the false positive rate (FPR) caused by the Tn5 bias toward open chromatins. The FPR is calculated by the number of peaks not overlapping with ChIP-seq but overlapping with ATAC-seq peaks, divided with the total number of peaks (**Fig. 3a**, **Supplementary Fig. 3**). The FPR for CoBATCH was as high as 46.8-54.3%, in contrast the FPR for CUT&Tag was 4.9-5.8%. For Stacc-seq, its FPR was 20.6-35.9%. The itChIP-seq and ULI-NChIP were almost not affected by open chromatin artefacts. Since the FPRs calculated here only considered the non-overlapping peaks, and because the overlapping peaks could also be generated from open chromatin, instead of real H3K27me3 peaks as exemplified in **Fig. 1b**, the FPRs presented here represented the lower limit. To calculate the FPR without a fixed p-value or q-value cutoff for the peaks, we assessed the FPRs for top peaks ranked by p-values for each method. Results shown in **Fig. 3b** indicated that most of the top peaks in CoBATCH represented open chromatin signals. Interestingly, while replicate 1 of the Stacc-seq showed lower FPR for the top peaks but the FPR gradually increased with more peaks, replicate 2 of Stacc-seq showed a relatively consistent FPR (**Fig. 3b**). Consistent with the high FPR, clustering analysis indicated that CoBATCH and Stacc-seq globally resembled open chromatin signals more closely than the H3K27me3 signals (**Fig. 3c**). These results indicate that CoBATCH and Stacc-seq have high false positive rates and thus great care should be taken in interpreting the data generated by these two methods.

**Fig. 3.**
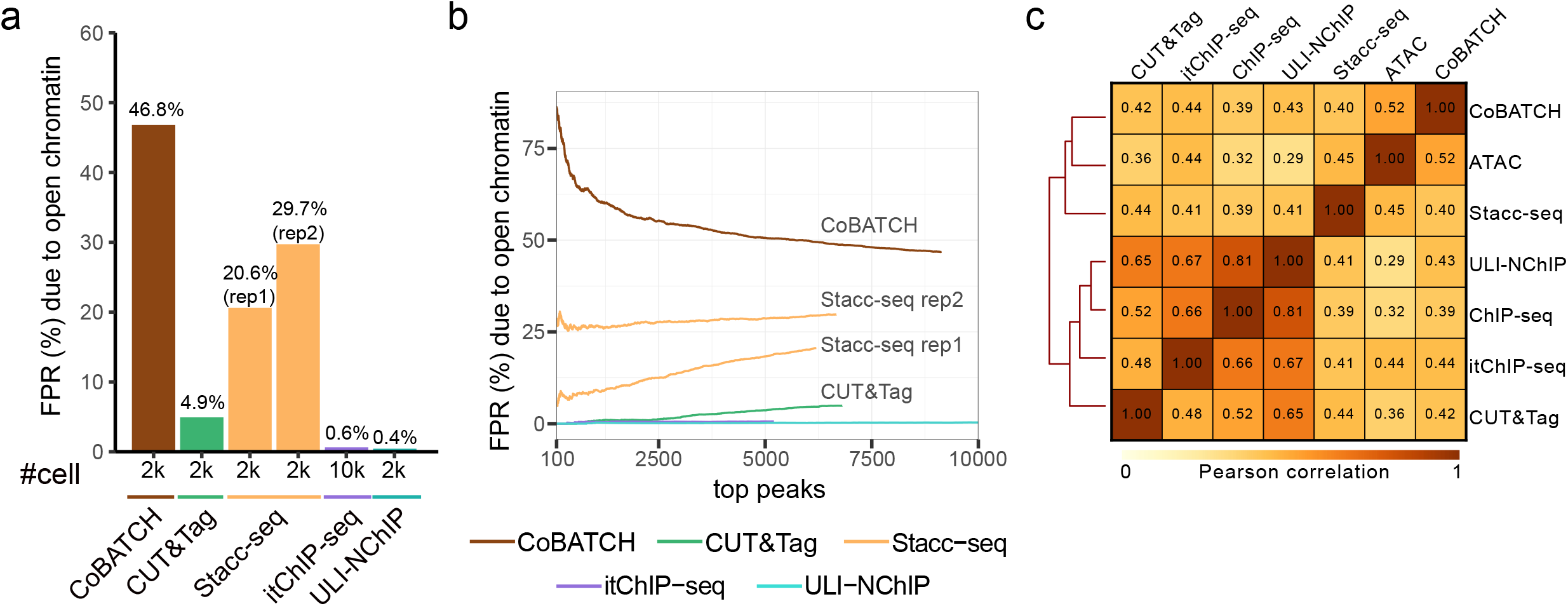
False positive rates of Tn5-based methods due to open chromatin. **a**, Overall false positive rate (FPR) due to open chromatin (measured by ATAC-seq) artefacts for each method. The number of cells (#cell) used for each library was indicated below each bar. **b**, False positive rate due to open chromatin (measured by ATAC-seq) artefacts for the top peaks in each method. **c**, Clustering of global H3K27me3 signals of each method with ATAC-seq and H3K27me3 ChIP-seq based on the Pearson correlation between any two methods (The Pearson correlation coefficients were shown in each box; bin size: 5kb).

## Discussion

In summary, our analysis reveals that the Tn5-based epigenomic profiling methods could capture substantial confounding open chromatin signals. The severity of open chromatin bias varies a lot among the different Tn5-based. CoBATCH and Stacc-seq are pone to open chromatin bias with high false positive rates. Thus, no matter adding the antibody and pA-Tn5 sequentially (CoBATCH) or pre-incubating antibody with pA-Tn5 and adding together (Stacc-seq), both procedures could result in high levels of bias toward open chromatin. Although CUT&Tag showed very weak H3K27me3 signals in non-PcG targets that resembled open chromatin signals, its overall false positive rate due to open chromatin is much lower than that from the CoBATCH despite both share almost identical experimental procedures (**Fig. 1a**). Thus, stringent washing with high salt before tagmentation employed in the CUT&Tag method must have contribute to the reduced open chromatin artefacts^23^, but would affect sites with weak binding. Signal normalization with IgG control could not eliminate the confounding signals from open chromatins for CoBATCH and Stacc-seq, while CUT&Tag without IgG control could have much lower biases. This coincides with the practice that no IgG control is needed for in situ immuno-cleavage-based profiling methods. The immunoprecipitation-based itChIP-seq utilizes Tn5 transposase to tagment DNA before immunoprecipitating the target DNA, thus its result is not affected by open chromatins. However, it is not an immunoprecipitation-free method, which requires input control, and its signal-to-noise intensities are not comparable to the immuno-cleavage-based methods when using small number of cells.

Given that Tn5-based methods are prone to open chromatin bias, cautions should be taken when the Tn5-based epigenomic profiling methods are used. We strongly recommend that evaluation of the confounding open chromatin signals and estimation of the FPR are performed under similar experimental conditions before these methods are used. We also suggest that in the future development of Tn5-based epigenomic profiling methods, repressive marks such as H3K27me3 or H3K9me3 should be used in evaluating the confounding open chromatin signals, instead of the active H3K4me3 mark used in the original publications of these methods. Since H3K4me3 largely colocalizes with open chromatins in mESCs, even the method mainly captures open chromatin signals, the use of H3K4me3 to evaluate would still show high correlation with bulk H3K4me3 ChIP-seq signals. Finally, cautions should be taken when interpreting data generated by Tn5-based epigenomic profiling methods due to the high FPR of open chromatin artefacts.

## Methods

### Data collection

In the original papers that described each of the Tn5-based method, most used H3K4me3 and/or H3K27me3 in mESCs for validation. However, the H3K4me3 in mESCs is mainly located at promoters in open chromatin regions. It is almost impossible to discriminate the peaks generated by true H3K4me3 or open chromatins. Thus, we used the repressed marker H3K27me3 in mESCs to evaluate the Tn5-based epigenomic profiling methods (summarized in Supplementary Table 1), which had the most publicly available datasets for different methods besides H3K4me3. For multiple replicates with the same or different number of cells for each method, we used the one with the best signal-to-noise ratio as the representative result for each method.

### Peak calling and signal track generation

For ChIP-seq, ULI-NChIP and Tn5-based methods, raw sequencing reads were first trimmed using Trimmomatic^27^ (version 0.39) to remove sequencing adaptors and low-quality reads. The cleaned reads were mapped to mm10 reference genome using bowtie2^28^ (version 2.4.2) with parameters: --local --very-sensitive-local --no-unal --no-mixed --no-discordant --dovetail -I 10 -X 700 --soft-clipped-unmapped-tlen. PCR duplicates were removed with Picard MarkDuplicates (version 2.23.4). Reads with mapping quality at least 30 were kept. For Tn5-based methods, proper paired reads with fragment length at least 178bp (nucleosome DNA size 140bp + 2 × Tn5 steric hindrance 19bp at both sides) were kept. For Tn5-based methods, to increase peaks resolution, the start and end positions for each fragment (one read pair) were shifted for 19bp toward internal to account for the steric hindrance of Tn5 enzyme. Peaks were called using MACS2^29^ callpeak (version 2.2.7.1) with parameters: -B – SPMR -p 1e-4 -g mm --broad --broad-cutoff 1e-4 --keep-dup all --scale-to large. Peaks were further filtered with q-value<0.01. The fold-change signal tracks were generated using MACS2 bdgcmp with input of treat-pileup and control-lambda bedgraph files generated from MACS2 callpeak in the last step. Peaks overlapping with mm10 blacklist regions (https://www.encodeproject.org/files/ENCFF547MET/) were removed. The ChIP-seq results were pooling of two replicates. The ULI-NChIP results were pooling of all four replicates. The ATAC-seq and DNase-seq datasets were analyzed using ENCODE ATAC-seq pipeline (version 1.9.3, https://github.com/ENCODE-DCC/atac-seq-pipeline).

### Peak comparison

Peaks were compared using bedtools^30^ intersect (version 2.29.2). Peaks with at least half-length intersecting with ChIP-seq / ATAC-seq / DNase-seq peaks were considered as overlapping (bedtools intersect parameters for getting overlapping peaks: -u -f 0.5; parameters for getting non-overlapping peaks: -v -f 0.5). The signal enrichment heatmaps were plotted using deeptools^31^ (version 3.5.0) computeMatrix and plotHeatmap. The TSSs of coding genes in the mouse genome were from GENCODE^32^ mouse gene set M24. The genome browser snapshot was generated with R package karyoploteR^33^ (version 1.18.0). The Pearson correlation and clustering analysis were performed using deeptools multiBigwigSummary and plotCorrelation with 5kb bin size and outliers removed.

## Data availability

The public datasets used in this study are summarized in Supplementary Table 1.

## Code availability

The code used to analyze the sequencing data is available at GitHub: https://github.com/YiZhang-lab/ChIPpipes

## Acknowledgements

We thank Drs. Chunxia Zhang and Zhiyuan Chen for discussion and critical reading of the manuscript. This project was supported by the NIH (R01HD092465) and the HHMI. Y.Z. is an Investigator of the Howard Hughes Medical Institute.

## Author contributions

Y.Z. supervised the project. M.W. performed the analysis. M.W. and Y.Z. wrote the manuscript.

## Supplementary for

**Supplementary Fig. 1.**
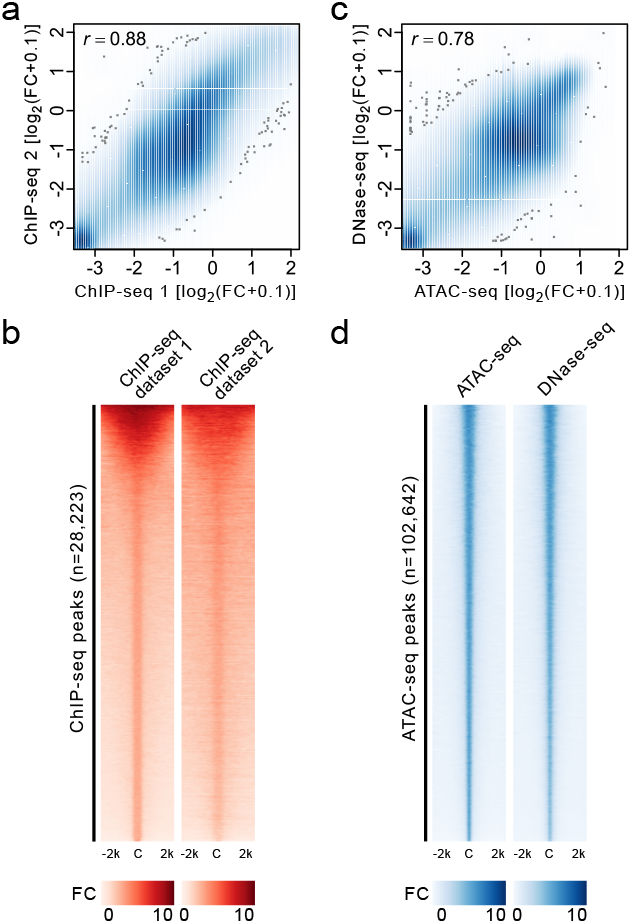
Different ChIP-seq and ATAC-seq datasets in mESC are consistent. **a**, Pearson correlation of two H3K27me3 ChIP-seq datasets in mESC (bin size: 5kb). **b**, Heatmap comparing H3K27me3 fold-change (FC) signals at the ChIP-seq peaks of two ChIP-seq datasets in mESCs (C: center of peaks in ChIP-seq dataset 1). **c**, Pearson correlation of ATAC-seq and DNase-seq in mESC (bin size: 5kb). **d**, Heatmap comparing open chromatin fold-change (FC) signals measured by ATAC-seq and DNase-seq in mESCs (C: center of peaks in ATAC-seq).

**Supplementary Fig. 2.**
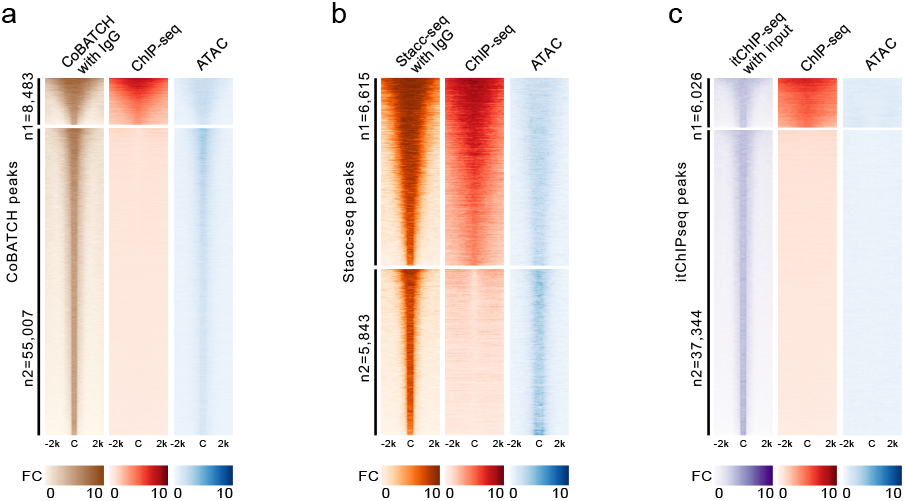
Evaluation of peaks called from different methods with input / IgG normalization. **a**, Significant peaks (q-value<0.01) called from CoBATCH with IgG control normalization were compared with open chromatin signals measured by ATAC-seq (n1: CoBATCH peaks overlapping with ChIP-seq peaks; n2: CoBATCH peaks not overlapping with ChIP-seq peaks; C: center of CoBATCH peaks; FC: fold-change over IgG control). **b**, Significant peaks (q-value<0.01) called from Stacc-seq with IgG control normalization were compared with open chromatin signals measured by ATAC-seq (n1: Stacc-seq peaks overlapping with ChIP-seq peaks; n2: Stacc-seq peaks not overlapping with ChIP-seq peaks; C: center of Stacc-seq peaks; FC: fold-change over IgG control). **c**, Significant peaks (q-value<0.01) called from itChIP-seq (100 cells) with input control normalization were compared with open chromatin signals measured by ATAC-seq (n1: itChIP-seq peaks overlapping with ChIP-seq peaks; n2: itChIP-seq peaks not overlapping with ChIP-seq peaks; C: center of itChIP-seq peaks; FC: fold-change over input control).

**Supplementary Fig. 3.**
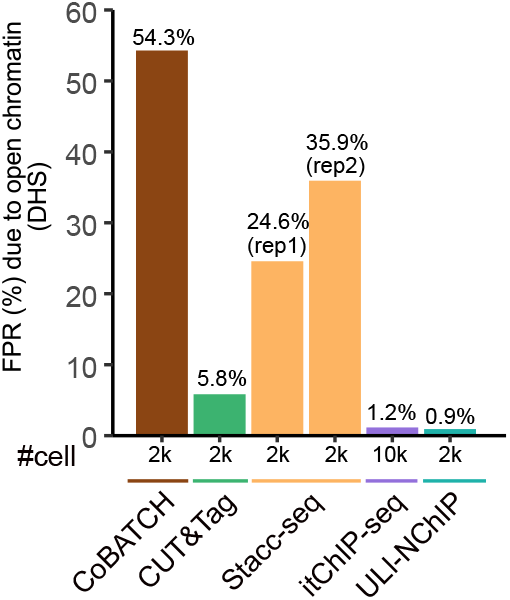
Overall false positive rate (FPR) due to open chromatin (measured by DNase-seq) artefacts for each method. The number of cells (#cell) used for each library was indicated below each bar.

**Supplementary Table 1.** Summary of public datasets used in this study.

| Method | Sample name | Cell number | Data source | Accession |
| --- | --- | --- | --- | --- |
| **CoBATCH** | 2k_mESC | 2,000 | GEO | GSM3711220 |
|  | 100_mESC_rep1 | 100 | GEO | GSM3711218 |
|  | 100_mESC_rep2 | 100 | GEO | GSM3711219 |
|  | IgG_2k_mESC_rep1 | 2,000 | GEO | GSM3893775 |
|  | IgG_2k_mESC_rep2 | 2,000 | GEO | GSM3893776 |
| **CUT&Tag** | 2k_mESC_rep1 | 2,000 | GEO | GSM4476407 |
|  | 2k_mESC_rep2 | 2,000 | GEO | GSM4476406 |
| **Stacc-seq** | 2k_mESC_rep1 | 2,000 | GEO | GSM4010607 |
|  | 2k_mESC_rep2 | 2,000 | GEO | GSM4010608 |
|  | IgG_mESC | NA | GEO | GSM4010609 |
| **itChIP-seq** | 10k_mESC_rep1 | 10,000 | GEO | GSM3609659 |
|  | 10k_mESC_rep2 | 10,000 | GEO | GSM3609660 |
|  | 100_mESC_rep1 | 100 | GEO | GSM3609661 |
|  | 100_mESC_rep2 | 100 | GEO | GSM3609662 |
|  | input | 10,000 | GEO | GSM3609658 |
| **ULI-NChIP** | 500_mESC_rep1 | 500 | GEO | GSM2082708 |
|  | 500_mESC_rep2 | 500 | GEO | GSM2082709 |
|  | 500_mESC_rep3 | 500 | GEO | GSM2082710 |
|  | 500_mESC_rep4 | 500 | GEO | GSM2082711 |
|  | input | 500 | GEO | GSM2082705 |
| **ChIP-seq**<br>**(dataset 1)** | mESC_rep1 | bulk | GEO | GSM2472743 |
|  | mESC_rep2 | bulk | GEO | GSM2472744 |
|  | input | bulk | GEO | GSM2472755 |
| **ChIP-seq**<br>**(dataset 2)** | mESC_rep1 | bulk | ENCODE | ENCFF001ZIB |
|  | mESC_rep2 | bulk | ENCODE | ENCFF001ZIH |
|  | input_rep1 | bulk | ENCODE | ENCFF001ZGK |
|  | input_rep2 | bulk | ENCODE | ENCFF001ZGM |
| **ATAC-seq** | 50k_mESC | 50,000 | GEO | GSM2156965 |
| **DNase-seq** | bulk | bulk | ENCODE | ENCSR000CMW |

